# GCTransNet: 3D mitochondrial instance segmentation based on Global Context Vision Transformers

**DOI:** 10.1101/2024.11.06.622389

**Authors:** Chaoyi Chen, Yidan Yan, Jingpeng Wu, Wen-Biao Gan

## Abstract

Mitochondria are double membrane-bound organelles essential for generating energy in eukaryotic cells. Mitochondria can be readily visualized in 3D using Volume Electron Microscopy (vEM), and accurate image segmentation is vital for quantitative analysis of mitochondrial morphology and function. To address the challenge of segmenting small mitochondrial compartments in vEM images, we propose an automated mitochondrial segmentation method called GCTransNet. This method employs grayscale migration technology to preprocess images, effectively reducing intensity distribution differences across EM images. By utilizing 3D Global Context Vision Transformers (GC-ViT) combined with global context self-attention modules and local self-attention modules, GCTransNet precisely models long-range and short-range spatial interactions. The long-range interactions enable the model to capture the global structural relationships within the mitochondrial segmentation network, while the short-range interactions refine local details and boundaries. In our approach, the encoder of the 3D U-Net network, a classical multi-scale learning architecture that retains high-resolution features through skip connections and combines multi-scale features for precise segmentation, is replaced by a 3D GC-ViT. The GC-ViT leverages shifted window-based self-attention, capturing long-range dependencies and offering improved segmentation accuracy compared to traditional U-Net encoders. In the MitoEM mitochondrial segmentation challenge, GCTransNet achieved state-of-the-art results, demonstrating its superiority in automated mitochondrial segmentation. The code and its documentation are publicly available at https://github.com/GanLab123/GCTransNet.

## 1. Introduction

Mitochondria, often referred to as the “powerhouses” of the cell, convert nutrients into adenosine triphosphate (ATP) through oxidative phosphorylation, providing essential energy for cellular functions. They possess a double-membrane structure: the outer membrane is smooth, while the inner membrane forms numerous cristae to increase surface area and enhance ATP production efficiency (Zick et al., 2009). In addition to its essential role in cellular metabolism (Rambold and Pearce, 2018), mitochondria are also involved in the processes of apoptosis (Polster and Pearce, 2004), calcium storage (Xu et al., 2016), and signal transduction (Campello and Scorrano, 2010; Cho et al., 2010). Mitochondrial dysfunction is closely associated with various diseases, including neurodegenerative diseases (Lin and Beal, 2006), metabolic disorders (Bhatti et al., 2017), cardiovascular diseases (Ong and Hausenloy, 2010), and certain types of cancer (Wallace, 2012).

Recent development of Volume Electron Microscopy (vEM) has revolutionized the field of microscopy, enabling researchers to obtain detailed three-dimensional views of biological samples (Peddie et al., 2022). In mitochondrial research, it has provided valuable insights into structure and function, and has the potential to advance our understanding of related diseases (Li et al., 2020).

In order to perform quantitative analysis and visualization of mitochondria, we need to segment individual mitochondria in the images. Traditional manual segmentation methods are subjective and time-consuming, whereas automated segmentation techniques, by incorporating Computer Vision and Machine Learning algorithms, can quickly and accurately segment mitochondria (Mekuč et al., 2020). This significantly saves researchers’ time and effort, reduces analytical errors, and ensures data reliability and consistency. Furthermore, automated segmentation techniques enable high-throughput analysis of large-scale mitochondrial image data, allowing researchers to rapidly extract meaningful information from vast datasets, and uncover potential patterns and trends. By automating mitochondrial segmentation, we can gain deeper insights into the complex relationships among mitochondrial morphology, cellular metabolism, energy production, and disease development (Bosch and Calvo, 2019).

Several mitochondrial electron microscopy datasets have been used in tasks such as segmentation, classification, and detection of mitochondria, aiming to help researchers better understand mitochondrial structure and function while reducing experimental workload. As early as 2011, Lucchi et al. published a small mitochondrial segmentation dataset known as the Lucchi dataset (Lucchi et al., 2011). Based on this dataset, they proposed a foundational segmentation model that significantly improved mitochondrial segmentation accuracy. Subsequently, the Lucchi team expanded this dataset, introducing the larger and more comprehensive Lucchi dataset. In 2015, Kasthuri et al. released a new mitochondrial segmentation dataset. Compared to Lucchi (Kasthuri et al., 2011), this dataset included a greater number of mitochondrial instances and larger data volume, which facilitated more challenging and diverse segmentation experiments with quantitiative evaluation. In 2020, Wei et al. released a large-scale mitochondrial instance segmentation dataset called the MitoEM dataset (Wei et al., 2020). This dataset included numerous structurally complex and morphologically diverse mitochondrial instances, greatly enriching the morphological data of mitochondria. Additionally, the MitoEM dataset contained numerous ultrastructures similar to mitochondrial structures, presenting new challenges for accurately segmenting mitochondria from complex backgrounds, prompting researchers to develop more advanced segmentation algorithms.

With the enhancement of computational power and the exponential growth of data, deep learning has made significant progress in various fields of Computer Vision. The introduction of FCN (Long et al., 2015) pushed segmentation technology to new heights. In 2015, Ronneberger et al. proposed the U-Net (Ronneberger et al., 2015) algorithm, which achieved multi-scale learning through skip connections. Due to its outstanding performance and generalization capability, U-Net has been widely applied in medical imaging, remote sensing, and data fusion. In the field of mitochondrial segmentation, researchers have proposed various improved network structures to address different challenges. For example, Xiao et al. (Xiao et al., 2018) proposed a fully residual 3D U-Net, which used 3D convolutional kernels and deep supervision, significantly detection task to capture mitochondrial shape priors. This method effectively reduced computational complexity and enhanced segmentation performance. Additionally, Luo et al. designed a multi-layer nested encoder-decoder structure to capture multi-scale contextual information (Luo et al., 2021). This fully utilized information from adjacent slices in a 2D segmentation network, achieving notable results.

In EM images, the similarity in grayscale and texture among various organelles brings significant interference to mitochondrial segmentation. To address this issue, Liu et al. improved the Mask-RCNN network, leveraging high-level semantic information to enhance the segmentation network’s classification ability, reducing interference from similar organelles in mitochondrial segmentation (Liu et al., 2020). Facing the challenge of large scale variations in mitochondrial instances under electron microscopy, Li et al. designed a simple yet effective anisotropic convolution block and deployed a multi-scale training strategy, thereby addressing this issue (Li et al., 2022). Pan et al. proposed an innovative Adaptive Template Transformer, which, through structural template learning and hierarchical attention mechanisms, effectively handled mitochondria of different scales and shapes (Pan et al., 2023).

Despite much efforts, previous 3D mitochondrial segmentation methods performed poorly in segmenting small targets, resulting in segmentation discontinuities where the same mitochondrion was incorrectly divided into different instances, as shown in **Fig. 1**. The first reason for this issue is the difficulty in feature extraction. The features of small targets may be similar to the surrounding environment, making it challenging to accurately distinguish them using traditional CNN feature extraction methods (Li et al., 2021). This leads to misjudgments in the segmentation model’s background around small targets, affecting segmentation accuracy and stability. Secondly, traditional CNN-based methods lack sufficient utilization of contextual information. In segmenting small targets, the contextual information of the surrounding environment is crucial for correctly locating and segmenting the target. Insufficient contextual information may prevent the model from accurately understanding the target’s position and shape.

**Fig. 1.**
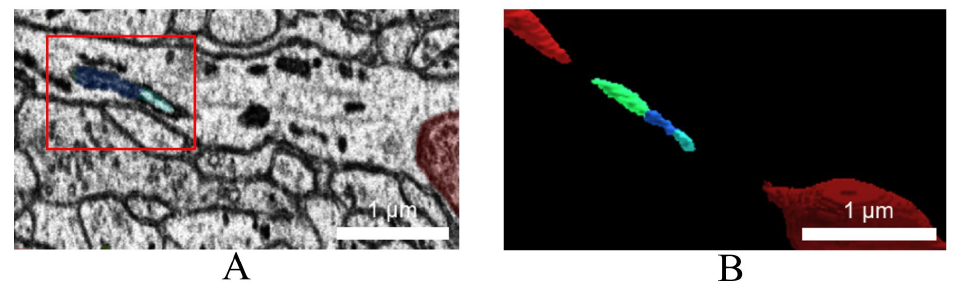
(A) Examples of Incorrect Segmentations by Previous Methods (Nightingale et al., 2021). (B) 3D Reconstruction of Erroneously Segmented Regions. Scale bar is 1 μm.

In this study, we developed a method to tackle this issue, and the primary contributions of our work are summarized as follows:

- We propose an electron microscopy image preprocessing method combining grayscale migration with median filtering to achieve normalization, thereby reducing grayscale differences between samples within the same dataset.
- We transform the Global Context Vision Transformers (Hatamizadeh et al., 2023) into a 3D form, fully utilizing 3D spatial contextual information. By integrating global context self-attention modules with standard local self-attention modules, we can effectively model both long-range and short-range spatial interactions.
- We replace the CNN-based encoder in the 3D U-Net network with a 3D GC-ViT encoder. The 3D GC-ViT leverages shifted window calculations for self-attention to extract features at different resolutions. These features are then connected to the 3D GC-ViT-based decoder at each resolution via skip connections.
- We demonstrate state-of-the-art performance by achieving top results in the MitoEM mitochondrial segmentation challenge (Franco-Barranco et al., 2022). Furthermore, we performed biological analysis on the segmentation results, quantifying the 3D morphology of mitochondria for comprehensive evaluation.

## 2. Material and Methods

### 2.1. Dataset

All experiments are based on a mitochondrial instance segmentation dataset constructed using the MitoEM dataset (Wei et al., 2020) and Lucchi++ dataset(Casser et al., 2020).

MitoEM dataset: This dataset utilizes multi-beam scanning electron microscopy to image two tissue blocks, generating two sub-datasets: MitoEM-H, containing mitochondrial data from the layer II of the human frontal cortex, and MitoEM-R, containing mitochondrial data from the layer II/III of the primary visual cortex of adult rats. Both tissue blocks have a volume of 30 cubic micrometers, ensuring high diversity in mitochondrial shapes and densities within the dataset. The MitoEM dataset comprises 500 human mitochondrial slices and 500 rat mitochondrial slices, each with a size of 4096 × 4096 pixels. The training set contains 400 human mitochondrial slices and 400 rat mitochondrial slices, while the validation set contains 100 human mitochondrial slices and 100 rat mitochondrial slices.

Lucchi++ dataset: This dataset, derived from the mouse hippocampus, includes both a training volume and a testing volume. Each volume has a resolution of 5 × 5 × 5 nm^3^ per voxel and is composed of 165 slices. The dimensions of each slice are 1024 × 768 pixels.

### 2.2. Pre-Processing

Regarding the various sources of noise interference in electron microscope images, such as electronic noise (Jones and Nellist, 2013) and speckle noise (Sim et al., 2004), median filtering is an effective nonlinear filtering method. It can significantly reduce these types of noise by replacing each pixel value with the median of the pixel values in its neighborhood, thereby eliminating outliers. This is highly beneficial for improving image quality and visualizing sample details. Unlike some linear filtering methods, median filtering (Huang et al., 1979) does not blur the image but instead preserves image details. This is particularly important for electron microscopic images, which typically contain rich microscopic structures and details that need to be retained for further analysis.

Since scanning electron microscope (SEM) (Mohammed and Abdullah, 2018) images are captured frame by frame, each image may exhibit variations in lighting or other imaging conditions. Grayscale consistency helps make semantic segmentation algorithms more robust to changes in illumination and imaging conditions. Even if the imaging conditions change or different imaging equipment is used, maintaining grayscale consistency allows the algorithm to accurately recognize semantic categories, enhancing its applicability and reliability in practical applications. However, common color transfer (Reinhard et al., 2001) methods are generally suitable only for three-channel color images and not for single-channel grayscale images, thus requiring some adjustments to the algorithms for grayscale image transfer, as shown in **Fig. 2(A)**.

**Fig. 2.**
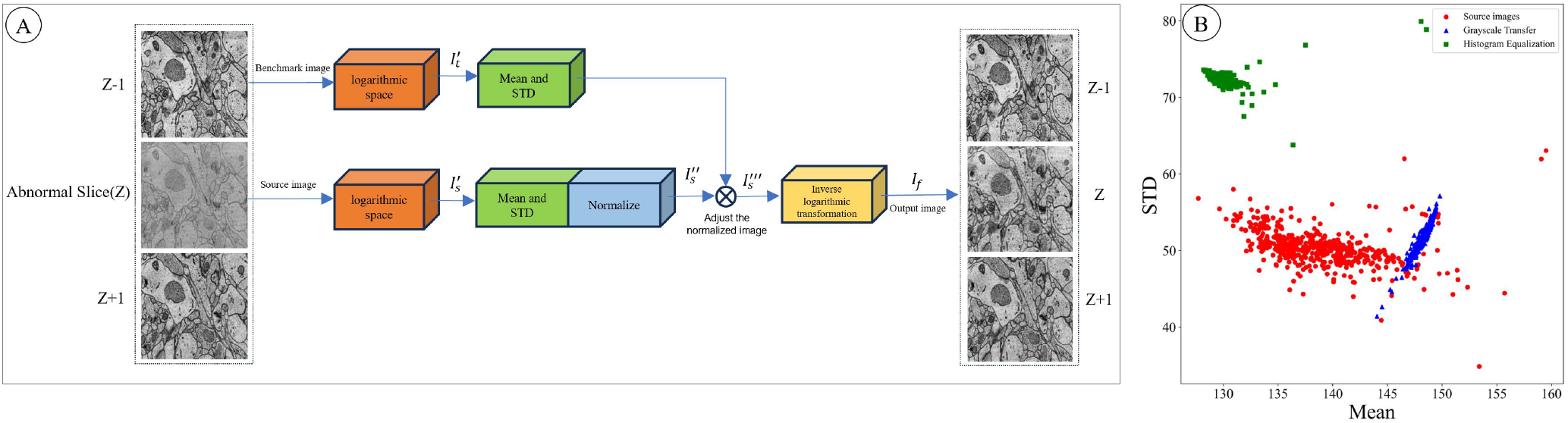
(A) Grayscale Transfer. The reference image and source image are first converted to logarithmic space, where their Means and STD are calculated. The source image is then normalized using these statistics to match the reference image’s characteristics. After adjustment, the normalized image is converted back to the original space to generate the final image. (B) Scatter plot of Mean and STD for Source images, Grayscale Transfer, and Histogram Equalization.

Firstly, compute the logarithmic values of the grayscale images 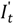 and 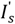. Logarithmic transformation can compress the high dynamic range of image data, making the brightness distribution more uniform and easier to process. Especially when there is a large variation in the brightness values of the image, the logarithmic space can effectively reduce the excessive expansion of the highlights while preserving the details in the darker regions. Suppose the target grayscale image is *I*_*t*_ and the source grayscale image is *I*_*s*_.

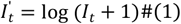

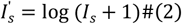

Next, calculate the Mean and Standard Deviation (STD) of the target and source images in the logarithmic space. The Mean and STD of the target image are μ_*t*_ and σ_*s*_, respectively. Normalize the source image with a Mean of 0 and STD of 1.

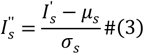

Adjust the Mean and STD of the standardized image to match those of the target image .

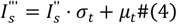

Perform the inverse logarithmic transformation on the adjusted source image to obtain the final image *I*_*f*_

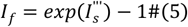

Through the above steps, Grayscale Transfer between grayscale images can be achieved. This process utilizes logarithmic transformation, adjustment of Mean and STD, and inverse logarithmic transformation to ensure fidelity of grayscale features in the transferred images.

**Fig. 2**(B) is a comparison between Grayscale Transfer and Histogram Equalization. Grayscale Transfer can achieve a concentration of the STD and Mean while approximating the original image style, and it results in fewer outliers compared to Histogram Equalization.

### 2.3. Modelstructureoverview

**Fig. 3(A)** illustrates the overall architecture of GCTransNet, which includes an encoder, bottleneck, decoder, and skip connections. In the encoder, mitochondrial electron microscope images are segmented into non-overlapping 4×4×4 patches, resulting in 64-dimensional features for each patch. These features are then projected into an arbitrary dimension through a linear embedding layer, passing through multiple GC-ViT blocks and patch merging layers to extract and integrate feature representations at different scales, capturing the hierarchical structure and semantic information of the images. The patch merging layers handle downsampling and dimensionality increase, while the GC-ViT blocks focus on feature representation learning.

**Fig. 3.**
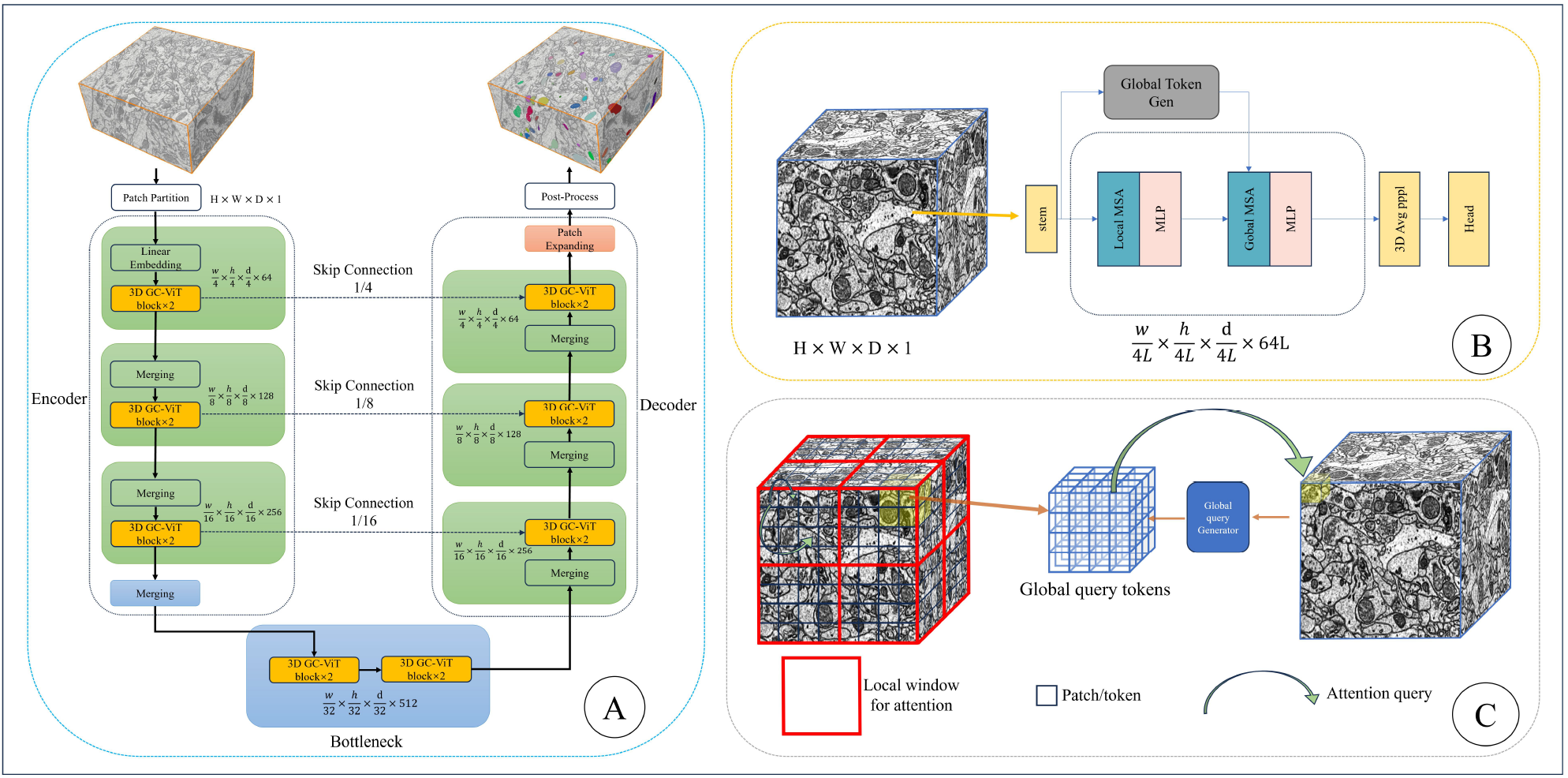
(A) The GCTransNet architecture. Our improvements are in the orange module in the figure. (B) The proposed 3D GC-ViT architecture. the query generator extracts 3D global query tokens, interacts with local keys and values to capture long-range information, and uses alternating 3D local and global context self-attention layers. (C) Attention mechanism forms. 3D global features are extracted from the input features and repeated to form 3D global query tokens, which interact with 3D local key and value tokens to capture long-range information.

Inspired by the design of U-Net, we devised a symmetric decoder based on 3D GC-ViT. The decoder performs a step-by-step upsampling process using 3D GC-ViT blocks and patch expanding layers. Through skip connections, high-resolution features in the decoder are fused with multiscale features from the encoder, effectively preserving details and spatial information in images, which is particularly crucial for applications like electron microscope images that require precise boundaries and structural details. The patch expanding layers are dedicated to upsampling, reshaping feature maps from adjacent dimensions into larger sizes. Finally, by applying 4x upsampling and a linear projection layer, pixel-level segmentation predictions are generated.

#### 2.4.. 3DGC-ViTblock

**Fig. 3(B)** depicts the architecture of 3D GC-ViT. We extended the GC-ViT model, originally designed for 2D image processing, to handle 3D data. 3D GC-ViT employs a hierarchical framework that progressively reduces spatial dimensions while expanding embedding dimensions to capture multiple resolution feature representations increased by powers of 2 (referred to as stages).

Initially, given an input image with resolution x, overlapping patches are obtained by applying a 3×3×3 convolutional layer with a stride of 2 and appropriate padding. These patches are then projected into a C-dimensional embedding space via another 3×3×3 convolutional layer.

Each stage of 3D GC-ViT consists of alternating local and global self-attention modules to extract spatial features. The local self-attention module operates within a local window similar to the Swin Transformer, while the global self-attention module accesses global features extracted by a global query generator, akin to a CNN-like module that extracts features of the entire image once per stage. At the end of each stage, spatial resolution is halved and channel count doubled through a downsampling block.

Inspired by the spatial feature contraction in CNN models, 3D GC-ViT introduces local biases and cross-channel interactions, effectively reducing dimensions. We utilize a modified 3D Fused-MBConv block followed by a max-pooling layer as the downsampling operator. The 3D Fused-MBConv can be represented by the following formula:

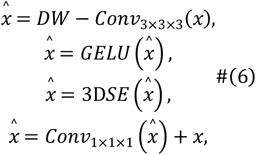

In the context provided: SE represents the 3D version of the Squeeze and Excitation module (Hu et al., 2018).GELU represents the Gaussian Error Linear Unit (Hendrycks and Gimpel, 2016).*DW*-*Conv*_×3×3_ represents a 3 × 3 × 3 depthwise convolution.

#### 2.5.. 3DGlobalself-Atention

**Fig. 3(C)** illustrates the form of attention. Local self-attention allows querying image patches within a local window, whereas global attention can query different image regions while still operating within the window. At each stage, the global query components are precomputed. Global self-attention utilizes extracted global query tokens shared across all blocks, interacting with local key and value representations. **Fig. 4** depicts the 3D Global query generator. The input feature map size is *H*× *W* × *D* ×*C*, where H denotes height, W denotes width, D denotes depth, and C denotes the number of channels. Initially, the input feature map undergoes a modified 3D Fused MBConv module for feature extraction and downsampling. Downsampling is achieved through max-pooling operations with a size of 2 × 2 × 2. This process is repeated 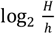 times to match the input feature size to the local window size h. The extracted feature map size becomes d×h×w×C, reshaped into a shape of (*d*·h ·*w*) × *C*. The reshaped feature map is replicated 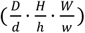 times, generating multiple local tokens. For the specific configuration of GCTransNet, please refer to Appendix A.

**Fig. 4.**
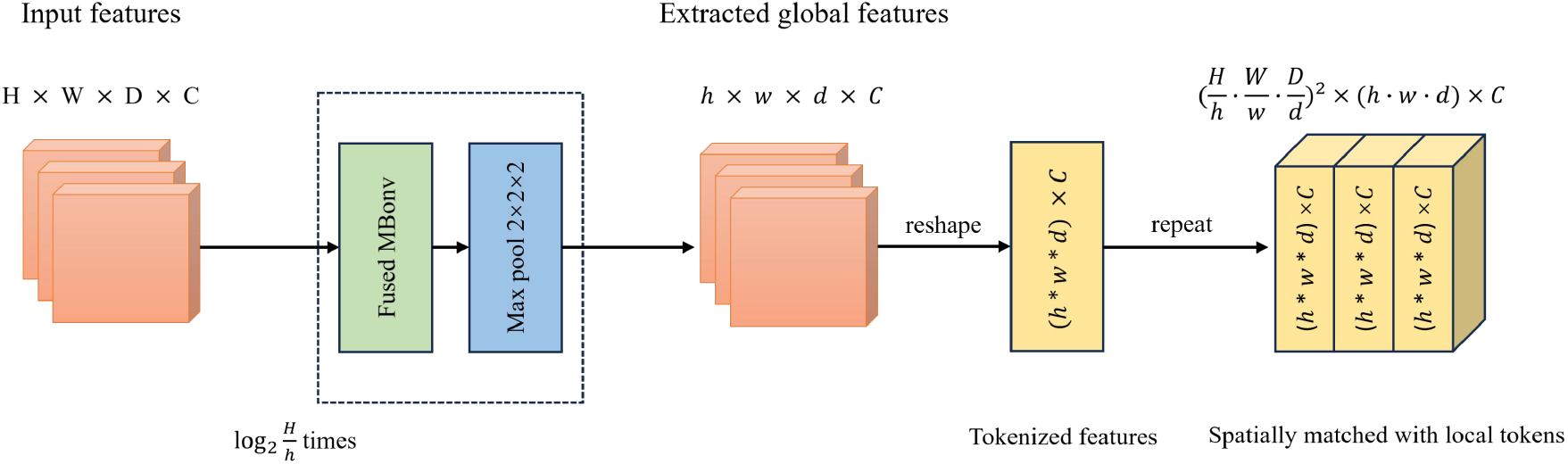
3D Global query generator. H, W, D, and C represent height, width, depth, and channel, respectively.

### 2.6.Post-Processing

In our study, we employ semantic segmentation models to segment images, categorizing each pixel into two classes: mitochondria (foreground) and non-mitochondria (background). However, semantic segmentation alone does not distinguish between different instances, necessitating post-processing of the binary segmented images. These algorithms include connected-component labeling (CC), morphological-based watershed (MBW), and marker-controlled watershed (MW). Our research utilizes morphological-based watershed as the post-processing method, adjusting it through operations like opening and closing to optimize the segmentation, followed by watershed transformation to separate clustered mitochondria

## 3.. Results

### 3.1. EvaluationMetricsandResults

Following the metrics of MitoEM (Wei et al., 2020) and Lucchi++ (Casser et al., 2020), we used Precision, Recall, Accuracy, and F1 score for quantitative evaluation. Precision measures the proportion of true positive samples among all samples predicted as positive by the model. The calculation formula is as follows:

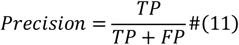

Recall is used to measure the model’s ability to correctly identify all samples that are truly positive (TP). The calculation formula is as follows:

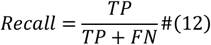

Where TP represents the number of samples correctly predicted as positive by the model, and FP represents the number of negative samples incorrectly predicted as positive by the model. FN represents the number of positive samples that were incorrectly predicted as negative by the model.

F1 score is the harmonic mean of Precision and Recall, used to comprehensively assess the model’s performance. The calculation formula is as follows:

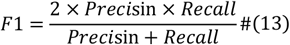

Accuracy in image segmentation refers to the ratio of correctly predicted pixels at the pixel level to the total number of pixels. In image segmentation tasks, images are typically divided into different regions or objects, and accuracy is an important metric to measure the performance of the model in such tasks.

Specifically, accuracy can be calculated using the following formula:

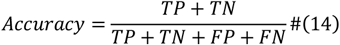

In this study, the performance of the proposed model was further evaluated through ablation experiments, using the following parameters:

**Method A**: 3D U-Net + CC

**Method B**: Grayscale Transfer + 3D U-Net + CC

**Method C**: Grayscale Transfer + 3D U-Net + CC + 3D GC-Vit

**Method D**: Grayscale Transfer + 3D U-Net + MBW + 3D GC-Vit

**Table 1** lists the parameter combinations used in different methods. **Table 2** presents the results of ablation experiments on the MitoEM validation set, specifically showing the Accuracy and F1 metrics for different methods on the MitoEM-R and MitoEM-H validation sets.

Method A is the baseline, lacking additional grayscale shift or 3D GC-Vit, and shows lower Accuracy and F1 scores on both datasets. Method B uses grayscale shift techniques, resulting in slight performance improvements, particularly on MitoEM-H. Method C integrates 3D GC-Vit, significantly enhancing all metrics, indicating its positive effect on performance. Method D builds on Method C by replacing post-processing with MBW, achieving the highest Accuracy and F1 scores, demonstrating MBW’s effectiveness in decoding 3D semantic segmentation results.

**Table 1.** Parameter combinations among different methods.

| Method | Grayscale Transfer | 3D GC-Vit | MBW |
| --- | --- | --- | --- |
| A |  |  |  |
| B | ✓ |  |  |
| C | ✓ | ✓ |  |
| D | ✓ | ✓ | ✓ |

**Table 2.**
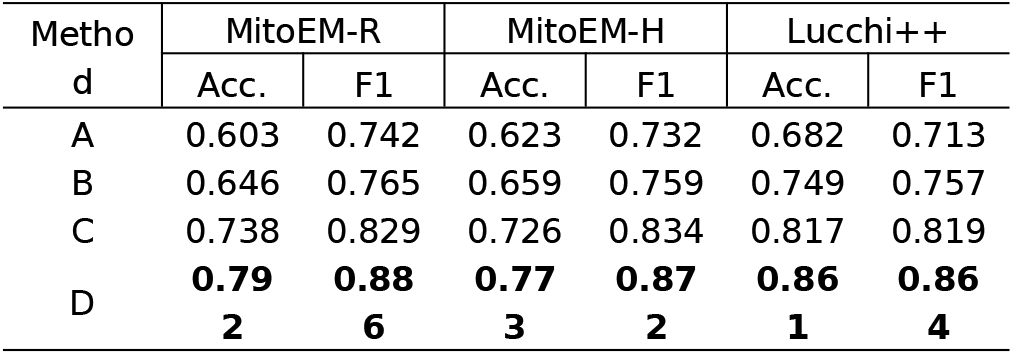
Comparative results of ablation experiments on the MitoEM validation set and Lucchi++ test set.

The compared methods are:

MitoNet (Glancy, 2023) : MitoNet is based on Panoptic-DeepLab (PDL) (Cheng et al., 2020) and employs a pre-trained ResNet50 as the encoder. It integrates two Atrous Spatial Pyramid Pooling decoders for semantic segmentation and instance prediction. The segmentation results are optimized using depthwise separable convolutions and the PointRend module.

VGG (Li et al., 2021): This method employs a voxel sampling strategy and a loss function with a similarity term to enhance voxel similarity and separability, using a 3D U-Net backbone for binary masks and boundary maps, and extracting instances with a watershed algorithm.

Res-UNet-R&H (Li et al., 2022): The Res-UNet-R and Res-UNet-H models segment 3D mitochondrial instances in electron microscope images, with Res-UNet-H featuring an enhanced decoder for noise reduction and an anisotropic convolution block for the MitoEM dataset.

FCI (Nightingale et al., 2021): Four convolutional neural networks based on a 5-level 3D U-Net were trained to predict mitochondrial binary masks and boundaries, using a smoothed Dice coefficient for loss and a watershed algorithm for instance extraction.

**Table 3** presents a comparative analysis of the proposed GCTransNet with U3D-BC (baseline) (Chen et al., 2016), MitoNet (Glancy, 2023), VGG (Li et al., 2021), Res-UNet-R&H (Li et al., 2022), and FCI (Nightingale et al., 2021). Some of the data are sourced from Franco-Barranco et al (Franco-Barranco et al., 2023). and the MitoEM mitochondrial segmentation challenge (Franco-Barranco et al., 2022). When comparing the performance of different methods on the MitoEM-R and MitoEM-H datasets, significant performance variations across different categories were observed. Overall, GCTransNet and Res-UNet-R&H exhibited the best performance on the Small category of the MitoEM-R dataset, while U3D-BC and GCTransNet demonstrated superior performance on the Medium category of the MitoEM-H dataset. Despite the generally lower F1 scores across methods in the Large category, performance in the Medium and Small categories was relatively better. Additionally, the overall performance on the MitoEM-R dataset was superior to that on the MitoEM-H dataset.

As illustrated in **Fig. 5**, the segmentation results of the GCTransNet model closely resemble the ground truth, featuring clear contours and complete shapes. Compared to other methods, it more accurately captures the boundaries of mitochondria, avoiding over-segmentation or under-segmentation. In the first row of images, where the mitochondrial targets are small, the GCTransNet model adapts well to these small targets, producing segmentation results highly consistent with the actual labels. In contrast, other models like Res-UNet-R&H and VGG perform poorly in handling these small targets, failing to capture all details accurately. In the fifth row of images, the GCTransNet model effectively preserves mitochondrial details and structures, whereas U3D-BC and FCI may produce blurred or incomplete results. GCTransNet excels in representing fine structures, ensuring high-quality segmentation outcomes. In the final row of images, the GCTransNet model accurately separates closely adhered mitochondria, with segmentation results closely matching the actual labels. Other models may exhibit unclear segmentation or incorrect connections when separating adhered mitochondria. GCTransNet’s finer feature extraction and segmentation algorithms allow it to more effectively identify and separate adhered mitochondria, ensuring the accuracy and completeness of the segmentation results.

**Table 3.** The comparative experimental results of GCTransNet with other methods on the MitoEM test set and Lucchi++ test set. The categories of mitochondria in the MitoEM dataset are small (length ≤ 1 μm), medium (1 μm < length < 4 μm), and large (length ≥ 4 μm).

| Method | Category | MitoEM-R |  |  |  | MitoEM-H |  |  |  | Lucchi++ |  |  |  |
| --- | --- | --- | --- | --- | --- | --- | --- | --- | --- | --- | --- | --- | --- |
|  |  | Prec. | Rec. | Acc. | F1 | Prec. | Rec. | Acc. | F1 | Prec. | Rec. | Acc. | F1 |
| U3D-BC<br>(Chen et al., 2016) | Large | 0.268 | 0.763 | 0.247 | 0.397 | 0.131 | **0.537** | 0.118 | 0.211 | 0.738 | 0.753 | 0.725 | 0.745 |
|  | Medium | 0.936 | 0.934 | 0.878 | 0.935 | 0.890 | **0.933** | 0.836 | 0.911 |  |  |  |  |
|  | Small | 0.479 | 0.910 | 0.457 | 0.628 | 0.662 | 0.917 | 0.625 | 0.769 |  |  |  |  |
| MitoNet<br>(Glancy, 2023) | Large | 0.207 | 0.498 | 0.171 | 0.292 | 0.167 | 0.439 | 0.137 | 0.242 | 0.713 | 0.752 | 0.709 | 0.732 |
|  | Medium | 0.822 | 0.831 | 0.705 | 0.826 | 0.827 | 0.841 | 0.715 | 0.834 |  |  |  |  |
|  | Small | 0.650 | 0.704 | 0.511 | 0.676 | 0.685 | 0.726 | 0.544 | 0.705 |  |  |  |  |
| VGG<br>(Li et al., 2021) | Large | 0.323 | 0.708 | 0.285 | 0.444 | 0.109 | 0.506 | 0.099 | 0.179 | 0.821 | 0.843 | 0.820 | 0.832 |
|  | Medium | 0.938 | 0.927 | 0.873 | 0.932 | 0.884 | 0.915 | 0.818 | 0.899 |  |  |  |  |
|  | Small | 0.494 | 0.919 | 0.473 | 0.643 | 0.603 | 0.922 | 0.574 | 0.729 |  |  |  |  |
| Res-U<br>Net-R&H<br>(Liet al., 2022) | Large | 0.214 | 0.515 | 0.178 | 0.302 | 0.117 | 0.341 | 0.096 | 0.174 | 0.839 | 0.837 | **0.858** | 0.838 |
|  | Medium | 0.353 | 0.794 | 0.323 | 0.489 | 0.172 | 0.549 | 0.151 | 0.262 |  |  |  |  |
|  | Small | **0.961** | **0.974** | **0.936** | **0.967** | 0.916 | **0.946** | **0.870** | **0.931** |  |  |  |  |
| FCI<br>(Nightingale et al., 2021) | Large | 0.362 | 0.229 | 0.540 | 0.281 | 0.167 | 0.470 | 0.141 | 0.246 | 0.822 | 0.835 | 0.815 | 0.828 |
|  | Medium | **0.969** | 0.833 | 0.813 | 0.896 | 0.821 | 0.831 | 0.703 | 0.826 |  |  |  |  |
|  | Small | 0.753 | 0.624 | 0.741 | 0.682 | 0.733 | 0.709 | 0.564 | 0.721 |  |  |  |  |
| GCTransNet<br>(ours) | Large | **0.612** | **0.823** | **0.541** | **0.702** | **0.194** | **0.537** | **0.166** | **0.285** | **0.842** | **0.887** | 0.853 | **0.864** |
|  | Medium | 0.967 | **0.954** | **0.923** | **0.960** | **0.923** | 0.923 | **0.857** | **0.923** |  |  |  |  |
|  | Small | 0.836 | 0.920 | 0.779 | 0.876 | **0.847** | 0.869 | 0.751 | 0.858 |  |  |  |  |

**Fig. 5.**
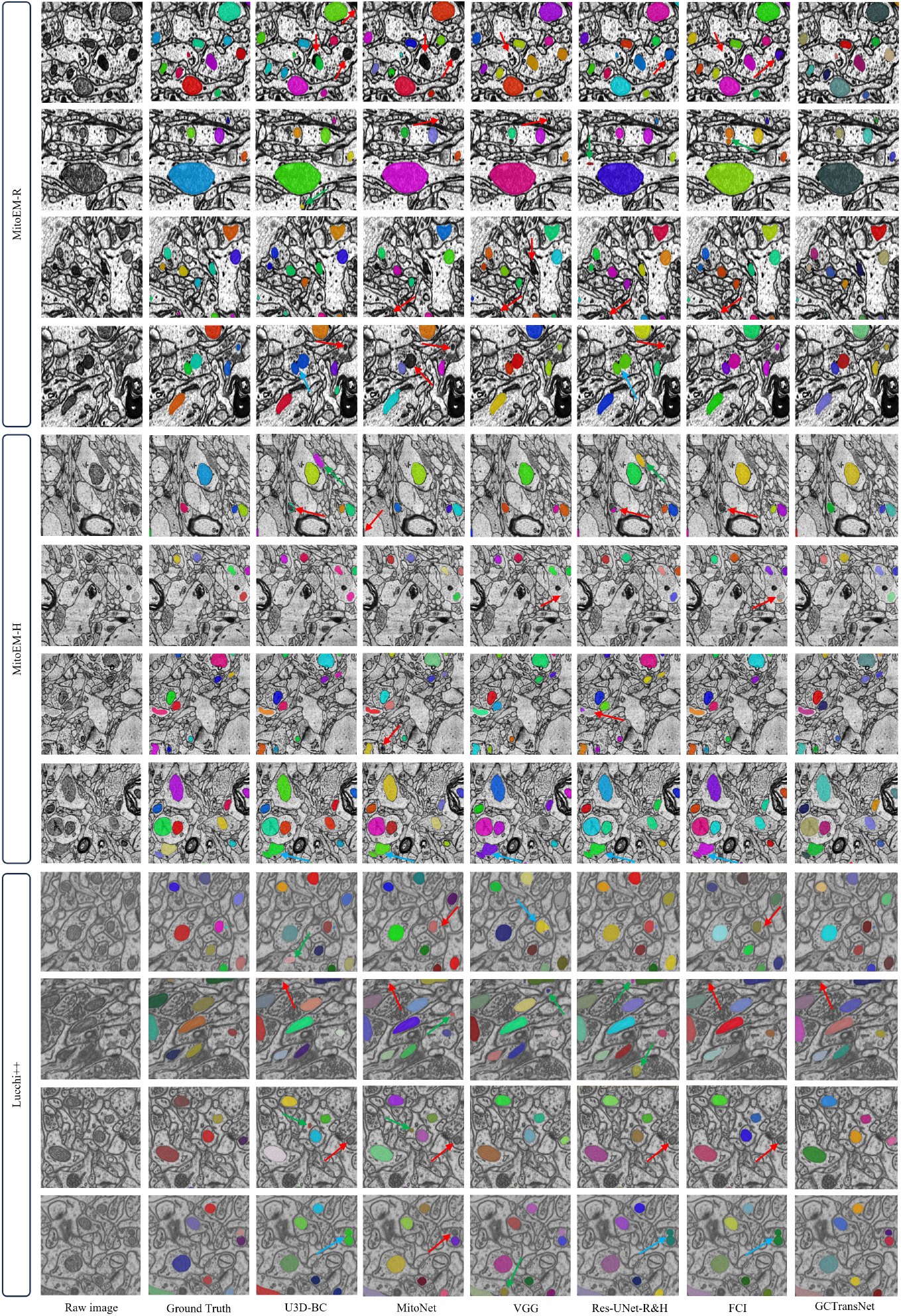
Comparison of GCTransNet Results with Other Models. The red arrow points to the missed segmentation area, the green arrow points to the positive segmentation area, and the blue arrow points to the merge error area.

**Fig. 6** illustrates the comparative 3D reconstruction results of GCTransNet and other models on the MitoEM-R, MitoEM-H, and Lucchi++ datasets. On the MitoEM-R dataset, GCTransNet achieves superior reconstruction with complete shapes and rich details, closely resembling the benchmark ground truth data. Other methods exhibit noticeable shape incompleteness and breaks. Similarly, on the MitoEM-H dataset, GCTransNet demonstrates excellent performance, producing smooth and complete reconstructions with high detail fidelity. However, due to the low signal-to-noise ratio and the presence of numerous artifacts in the MitoEM-H dataset, small targets still experience breaks. Due to the relatively simple mitochondrial shapes in the Lucchi++ dataset, there is no significant difference in the results between GCTransNet and the other models. Overall, GCTransNet’s reconstruction results on both datasets surpass those of other models, showcasing its higher reconstruction integrity and detail restoration capability.

**Fig. 6.**
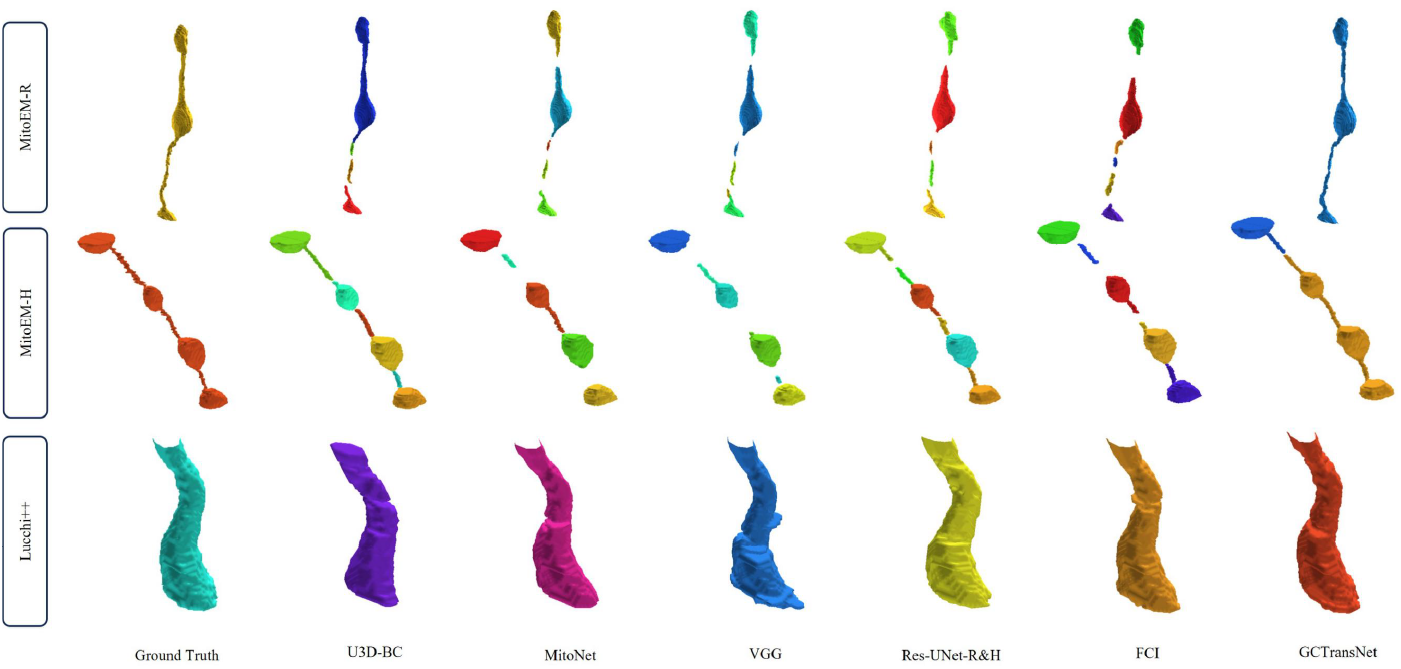
Comparison of 3D Reconstruction Results Between GCTransNet and Other Models.

#### 3.2.. 3DMorphologyofMitochondria

Mitochondria are organelles within cells responsible for producing the energy currency of the cell, adenosine triphosphate (ATP). The morphology of mitochondria is closely related to their function, making the study of their three-dimensional morphology crucial for understanding their biological roles.

In cells, mitochondrial morphology can vary due to cell type, cellular state, and environmental conditions. As shown in Fig. **7**, the MitoEM dataset reveals various morphologies of mitochondria, including elongated tubular shapes, reticular networks, and spherical forms. These morphological variations may correlate with cellular energy demands, metabolic states, and physiological processes such as apoptosis.

**Fig. 7.**
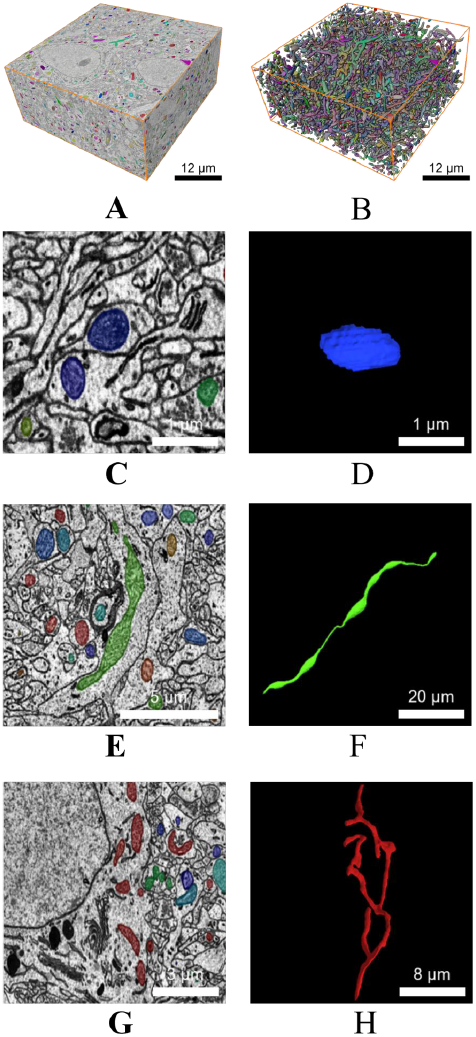
The 3D morphology of mitochondria at different locations within neurons. (A) is the 3D instance segmentation result, and (B) is the 3D reconstruction result; Scale bar is 12 μm. (C, D) Mitochondria (blue) are located at synaptic sites; Scale bar is 1 μm.(E, F) Mitochondria (green) are situated in axons and dendrites; Scale bar is 5 μm (E), 20 μm (F).(G, H) Mitochondria (red) are found in the cell body; Scale bar is 3 μm (G), 8 μm (H).

Different shapes of mitochondria at different locations may reflect their varied functional and metabolic demands within the cell. At synaptic sites, mitochondria are suggested to regulate synaptic activity through energy modulation. Given the rapid energy requirements at synapses for neurotransmitter release and other activities, synaptic mitochondria often exhibit flattened shapes to efficiently supply energy.

In axons and dendrites, mitochondrial networks adapt into elongated tubular structures surrounding microtubules. This morphology facilitates ATP provision along axonal or dendritic lengths, supporting the extensive energy demands of axonal and dendritic extensions.

Within the cell body, mitochondria typically form a reticular network radiating from the nucleus. This networked morphology helps mitochondria support overall cellular energy needs and facilitates the transport of energy and metabolic products among different organelles. Mitochondria in the cell body must meet the overall energy demands of the cell and efficiently transfer energy and metabolites between different organelles, thus forming an interconnected network to accomplish these functions effectively.

### 3.3.. MeasurementofMitochondrialMorphology

When measuring the morphological parameters of 3D mitochondria, several key parameters are particularly noteworthy as they provide important information about mitochondrial structure and function. These parameters include:

#### Volume and Surface Area

The volume of mitochondria refers to the amount of space they occupy. Mitochondrial volume can reflect their potential for energy production and metabolic activity within cells. Abnormal mitochondrial volume may be associated with cellular dysfunction. Mitochondrial surface area refers to the total area of their outer surface. Surface area is related to mitochondrial respiration and metabolic activities, as the outer surface is involved in exchanges with other molecules in the cytoplasm.

#### Shape Factor

The shape factor of mitochondria describes characteristics of their shape, often measured by parameters such as the ratio of mitochondrial length, width, and thickness. Shape factors may be associated with the functional state and metabolic activity of mitochondria.

We utilized Amira software to statistically analyze the MitoEM dataset for mitochondrial Number, Density, Surface Area, and Volume, as shown in **Table 4**. Overall, human mitochondria show significantly higher numbers and density compared to rats. Specifically, human mitochondria number 9294, whereas rats have 5904. Human mitochondrial density is 0.344 N/μm^3^, while rats have 0.219 N/μm^3^. However, rat mitochondria exhibit significantly larger surface areas and volumes than humans, measuring 2.318 μm^2^ and 0.216 μm^3^ respectively, compared to 0.982 μm^2^ and 0.075 μm^3^ for humans. Additionally, the surface area to volume ratio of human mitochondria (17.478) is higher than that of rats (14.705). This suggests that human mitochondria have more complex surface features, potentially reflecting their ability to efficiently exchange substances and convert energy to adapt to different cellular environments and functional needs. In contrast, the 3D Sphericity of rat mitochondria (0.767) is lower than that of humans (0.775), indicating that the shape of rat mitochondria is more irregular or complex. This could be related to the high dynamic variability of mitochondria, such as frequent fission and fusion processes. The 3D Sphericity is a quantitative measure used to describe the compactness of an object, defined as:

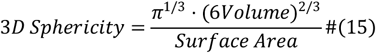

**Table 4.** Statistical data of mitochondria in the MitoEM dataset, including Number, Density(N/μm^3^), Surface Area(μm^2^), Volume(μm^3^), Surface Area/Volume, and 3D Sphericity.

| Dataset | Category | Number | Density | Surface Area | Volume | Surface Area/Volume | 3D Sphericity |
| --- | --- | --- | --- | --- | --- | --- | --- |
| MitoEM-R | All | 5904 | 0.219 | 2.318 | 0.216 | 14.705 | 0.767 |
|  | Small | 2656 | 0.098 | 0.771 | 0.055 | 17.178 | 0.862 |
|  | Medium | 2879 | 0.106 | 2.189 | 0.19 | 13.091 | 0.715 |
|  | Large | 37 | 0.013 | 14.449 | 1.581 | 9.518 | 0.477 |
| MitoEM-H | All | 9294 | 0.344 | 0.982 | 0.075 | 17.478 | 0.775 |
|  | Small | 6946 | 0.257 | 0.659 | 0.045 | 18.576 | 0.873 |
|  | Medium | 2311 | 0.085 | 1.856 | 0.156 | 14.273 | 0.719 |
|  | Large | 37 | 0.001 | 7.164 | 0.68 | 11.348 | 0.525 |

The larger the 3*D Shericity* the less compact the object’s shape, potentially exhibiting a more complex morphology.

As shown in Fig. **8**, these data collectively reveal significant differences between rat and human mitochondria in terms of number, size, density, and shape. This suggests that there may be distinct adaptive features in structure and function between the two species.

**Fig. 8.**
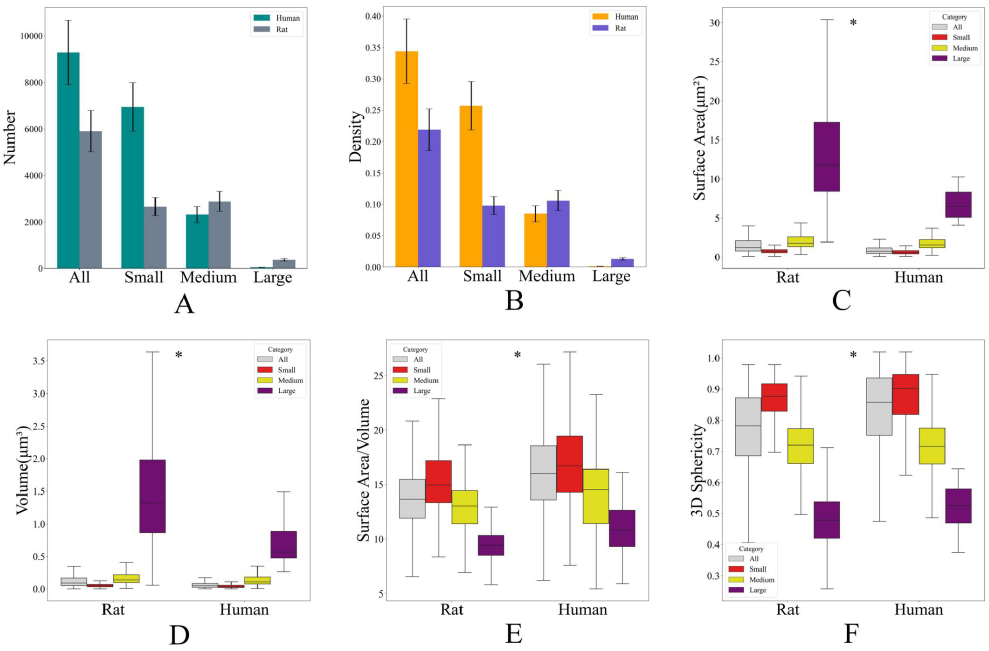
Statistics of mitochondria in the MitoEM test set. (A) Measurement of quantity. (B) Measurement of density. (C) Measurement of surface area. (D) Measurement of volume. (E) Measurement of the ratio of surface area to volume. (F) Measurement of the 3D Sphericity. \**P*≤ 0.01.

**Table 5** presents the statistical data on Length, Breadth, Thickness, and their respective ratios within the MitoEM test dataset. As shown in **Fig. 9**, there are significant differences between rat and human mitochondria in terms of length, breadth, thickness, and their ratios. The average length of rat mitochondria is 1.667 μm, the breadth is 0.613 μm, and the thickness is 0.418 μm. These values are higher than those of human mitochondria, which have an average length of 0.850 μm, a breadth of 0.513 μm, and a thickness of 0.345 μm. Additionally, the ratios of the rat mitochondria are higher than those of humans, with a length-to-breadth ratio of 2.486 compared to 1.659, and a length-to-thickness ratio of 3.622 compared to 2.525, indicating that rat mitochondria are longer and more elongated. In contrast, human mitochondria are relatively more spherical, with more balanced proportions. The breadth-to-thickness ratios are quite similar, with the rat at 1.489 and the human at 1.531, showing little difference between the two in this regard.

**Table 5.** The statistical data on Length(μm), Breadth (μm), Thickness (μm), and their respective ratios within the MitoEM test dataset.

| Dataset | Category | Length | Breadth | Thickness | Length/Breadth | Length/Thickness | Breadth/Thickness |
| --- | --- | --- | --- | --- | --- | --- | --- |
| MitoEM-R | All | 1.667 | 0.613 | 0.418 | 2.486 | 3.622 | 1.489 |
|  | Small | 0.713 | 0.476 | 0.347 | 1.555 | 2.224 | 1.436 |
|  | Medium | 1.607 | 0.647 | 0.433 | 2.629 | 3.867 | 1.517 |
|  | Large | 9.001 | 1.336 | 0.787 | 8.059 | 11.772 | 1.652 |
| MitoEM-H | All | 0.850 | 0.513 | 0.345 | 1.659 | 2.525 | 1.531 |
|  | Small | 0.625 | 0.463 | 0.32 | 1.392 | 2.078 | 1.496 |
|  | Medium | 1.454 | 0.657 | 0.418 | 2.393 | 3.755 | 1.635 |
|  | Large | 5.359 | 1.034 | 0.595 | 5.853 | 9.645 | 1.769 |

**Fig. 9.**
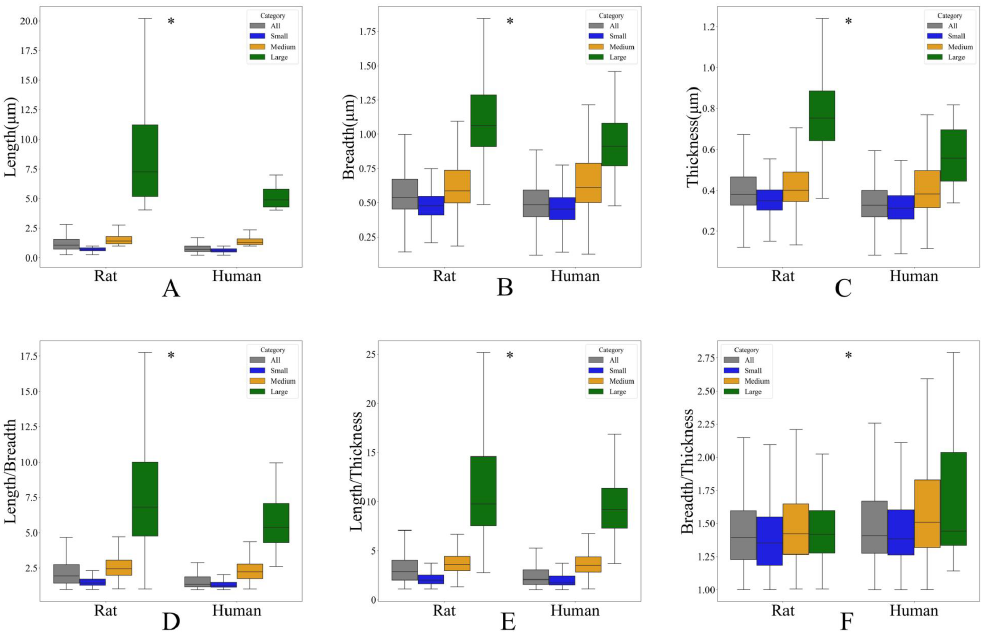
Morphological measurements of mitochondria in the MitoEM test set. (A) Length (B) Breadth (C) Thickness (D) Length/Breadth (E) Length/Thickness (F) Breadth/Thickness. \**P*≤ 0.01.

## 4. Conclusion

In this study, we propose GCTransNet, a 3D mitochondrial instance segmentation model based on Global Context Vision Transformers. This approach employs Grayscale Transfer techniques to preprocess images, significantly reducing grayscale variations among different slices. By integrating 3D Global Context Vision Transformers, the method leverages global and local self-attention modules to accurately model long-range and short-range spatial interactions. The performance of the model is enhanced by incorporating the GC-ViT encoder into the 3D U-Net network and using the shifted window mechanism to compute self-attention.

When compared with existing methods, our approach excels on the MitoEM test set. In the large category of the MitoEM-R subset, GCTransNet achieves an Accuracy of 0.541 and an F1 score of 0.702, significantly surpassing other methods. In the medium category of the MitoEM-R subset, GCTransNet attains an Accuracy of 0.923 and an F1 score of 0.960, slightly higher than the top competitor. In the medium category of the MitoEM-H subset, GCTransNet achieves an Accuracy of 0.857 and an F1 score of 0.923, markedly outperforming other methods. These results demonstrate that GCTransNet exhibits superior overall performance in the task of 3D mitochondrial instance segmentation in volumetric electron microscopy, highlighting its potential and efficacy in biomedical image analysis.

## Code and Data availability

The code is publicly available at https://github.com/GanLab123/GCTransNet, and the datasets are publicly available at https://mitoem.grand-challenge.org and https://casser.io/connectomics/.

## Acknowledgements

This work was supported by Lingang Laboratory, the National Key R&D Program of China (No.2022YEF0203200) and the STI2030-Major Project (No. 2022ZD0211900).

## Appendix A.. An example appendix

### A.1.. GCTransNetconfigurations

- Embed Dimension: 768
- Feature Size: 48
- Number of Blocks: [2,2,2,2]
- Window Size: [7,7,7]
- Number of Heads: [3,6,12,24]
- Parameters: 49.24M
- FLOPs: 465.28G

